# Ultrastructural Analysis of Human Uncinate Fasciculus with Spectral-Focusing Coherent Anti-Stokes Raman Spectroscopy

**DOI:** 10.1101/2025.09.18.672977

**Authors:** Kelly Perlman, Justine Major, Armand Collin, Valérie Pineau Noël, Murielle Mardenli, Sébastien Jerczynski, Maria Antonietta Davoli, Julien Cohen-Adad, Daniel Côté, Naguib Mechawar

**Author notes:** these authors contributed equally (Kelly Perlman, Justine Major, Armand Collin).

## Abstract

Characterizing the ultrastructure of myelin in the human brain is key to understanding the neurobiology of both health and disease. In postmortem human brain tissue, electron microscopy is often technically unfeasible due to poorer tissue quality. The uncinate fasciculus (UF) is a long-range white matter association tract that connects the anterior temporal lobe with the orbitofrontal cortex. The UF is not present in rodents yet is highly expanded in humans and non-human primates. As such, its molecular and ultrastructural properties are virtually unknown. Here, we develop and validate a novel spectral-focusing Coherent Anti-Stokes Raman Spectroscopy (sf-CARS) system coupled with a custom AxonDeepSeg segmentation model to characterize UF ultrastructure in the human postmortem brain (n=6). We provide a proof of concept of this new methodological pipeline in the UF temporal segment and observe that the mean axon diameter detected is 0.93 µm ± 0.54 and mean myelin thickness is 0.48 µm ± 0.14. We also observe that the UF axons are thicker than those in the anterior cingulate cortex white matter. We detail and validate the full methodology, including tissue fixation and sectioning, sf-CARS acquisition settings, as well as the AxonDeepSeg deep learning model parameters such that this pipeline can be utilized by others in the field.

## Introduction

The uncinate fasciculus (UF) is a hook-shaped association white matter fiber tract that bidirectionally connects the anterior temporal lobe to the medial and lateral orbitofrontal cortex and part of the inferior frontal gyrus (Liakos et al., 2021; Von Der Heide et al., 2013). This tract is sometimes referred to in segments, namely the temporal segment, the insular segment, and the frontal segment (Peltier et al., 2010; Von Der Heide et al., 2013). This tract is not present in rodents (not to be confused with the uncinate fascicle of the rodent cerebellum (Allen Institute for Brain Science, 2011; Liang et al., 2017)), and has been studied mostly in humans and non-human primates via diffusion tensor imaging or gross dissection studies, and as such it is virtually uncharacterized from molecular and ultrastructural perspectives (Von Der Heide et al., 2013). The closest information available is from a study that used electron microscopy to look at the point of intersection between the UF and the inferior occipitofrontal fasciculus in 3 human brains donated for medical education, with 4% formaldehyde fixation via infusion through the femoral artery less than 48 hours postmortem (Liewald et al., 2014). Electron microscopy in postmortem human brains is possible provided that preservation conditions are excellent (Sele et al., 2019) such as those in rapid autopsy programs, however, much of the postmortem human tissue available has longer postmortem intervals (PMI), and is in poorer condition. As such, electron microscopy is unfeasible for most collected postmortem brain tissue. This is especially the case for brain banks with specialized focus like suicide, in which the brains typically have longer PMIs and refrigeration delays (time between death and storage of the body at 4°C), like those of the Douglas Bell-Canada Brain Bank (“Brain Donation Procedures in the Sudden Death Brain Bank in Edinburgh,” 2018).

Coherent Anti-Stokes Raman Scattering (CARS) microscopy presents an alternative method to obtain ultrastructural metrics, with a high sensitivity, label-free technique (Bégin et al., 2013, 2014; Bélanger et al., 2010; Cheng & Xie, 2013; Turcotte et al., 2016). This method capitalizes on the strong Raman scattering properties of lipids, by probing the methylene (CH_2_) stretching vibrational band (2,845 cm^-1^) (Evans et al., 2005; Lutz et al., 2017). Fatty acid tails are rich in CH_2_ groups, and since myelin is extremely lipid rich (approximately 70-85% lipids by dry weight (Poitelon et al., 2020)) using the CH_2_ vibrational signature essentially creates a molecular map of myelin in a tissue slice.

Previously, our group utilized CARS microscopy to study the postmortem anterior cingulate cortex (ACC) in the context of depression and childhood abuse (CA) (Lutz et al., 2017). We observed alterations in gene expression and DNA methylation patterns related to myelination as well as ultrastructure changes exclusively in depressed individuals who died by way of suicide with a history of severe CA (Lutz et al., 2017). Specifically we found that, in small caliber axons (0.5-1.25 µm diameter) of the ACC white matter, individuals with a history of CA had thinner myelin and larger g-ratios than the comparator groups (Lutz et al., 2017).

Since the publication of these ACC results, a newer version of CARS microscopy has been developed. Spectral-focusing CARS (sf-CARS) represents a significant advancement in label-free neurophotonics imaging, overcoming key limitations of traditional CARS systems that rely on picosecond pulses. By employing ultra-short femtosecond pulses across larger wavebands, sf-CARS utilizes temporal chirping through a glass rod to dramatically enhance spectral resolution and image contrast (Pegoraro et al., 2009). This innovative approach enables precise targeting of different Raman bands through controlled temporal overlap between pulses, facilitating fast hyperspectral imaging via automated delay stage manipulation. While the absorption cross-section of myelin is not naturally optimized for femtosecond pulse interactions, the chirping process creates a non-linear dispersion over time of frequencies within the envelope of the pulse, and with it, a reduced effective power per spectral component. This approach still enables successful myelin targeting despite the power-dependent nature of Raman scattering.

Furthermore, since the publication of our ACC results, segmentation approaches have been augmented with the implementation of deep learning. While manual quantification remains the gold standard for morphometric assessment (Ma et al., 2024), this approach is labor-intensive, requiring extensive expert annotation time and specialized histological expertise. Recent advances in deep learning have introduced more efficient alternatives, particularly the U-Net architecture (“U-Net,” 2015) which has demonstrated superior time efficiency and practicality compared to traditional manual methods for biomedical image segmentation. AxonDeepSeg represents a state-of-the-art segmentation framework that employs a U-Net-based architecture to perform multi-class semantic segmentation of axons and myelin sheaths while simultaneously extracting comprehensive morphometric parameters (Zaimi et al., 2018). This approach based on convolutional neural networks offers several advantages over manual quantification: reduced annotation burden for large-scale datasets, decreased processing time, and automated extraction of key morphometric features. The framework can effectively leverage manually annotated training data to generate accurate predictions on custom datasets across diverse microscopy modalities.

Therefore, with the advances in both microscopy and image segmentation technologies, we provide an updated, modernized workflow to quantify white matter ultrastructure in the postmortem human brain. Here, we validate this new method on the UF, providing both a proof of concept and the first ever ultrastructural metrics of the UF temporal segment.

## Methods

### Tissue collection

This research was approved by the Douglas Hospital Research Ethics Board. Frozen UF was dissected from the left Brodmann Area 38 (temporal pole). It was dissected from this region because the WM tract is purely UF and not contaminated with adjacent fibers such as the inferior frontal occipital fasciculus. The brains were obtained through the Douglas-Bell Canada Brain Bank (www.douglasbrainbank.ca), thanks to a close collaboration with the Québec coroner’s office. Informed consent was obtained from the donors’ next of kin, and donor phenotypic information was acquired by way of a standardized psychological autopsy procedure. In this proof-of-concept study, we excluded subjects with psychiatric disorders and histories of childhood abuse and focused on “control” subjects only. Neurodegenerative disorders also constituted an exclusion criterion.

### Tissue fixation and sectioning

Blocks of fresh frozen UF (approximately 1 cm^3^) were fixed overnight in 10% formalin at 4°C and then stored in 1x PBS. The fixed tissue was then cut on a Leica VT1200S vibratome filled with ice cold 1x PBS at 300 µm thick sections (settings: - 0.8 mm amplitude, 0.8 mm/s speed, 300 µm step size) (Figure 1A). The slices were stored in vials with PBS with cryopreservative (glycerol and ethylene glycol) at -20 °C. Since the original orientation of the tissue at dissection was not preserved in the tissue cassettes, each block was split into 3 and sliced in all 3 orientations with respect to a preselected face of the cube. Upon imaging, the one section with the clear, abundant, dense fibers was selected as the correct orientation. The imaging was performed at the CERVO brain research center at Université Laval.

**Figure 1.**
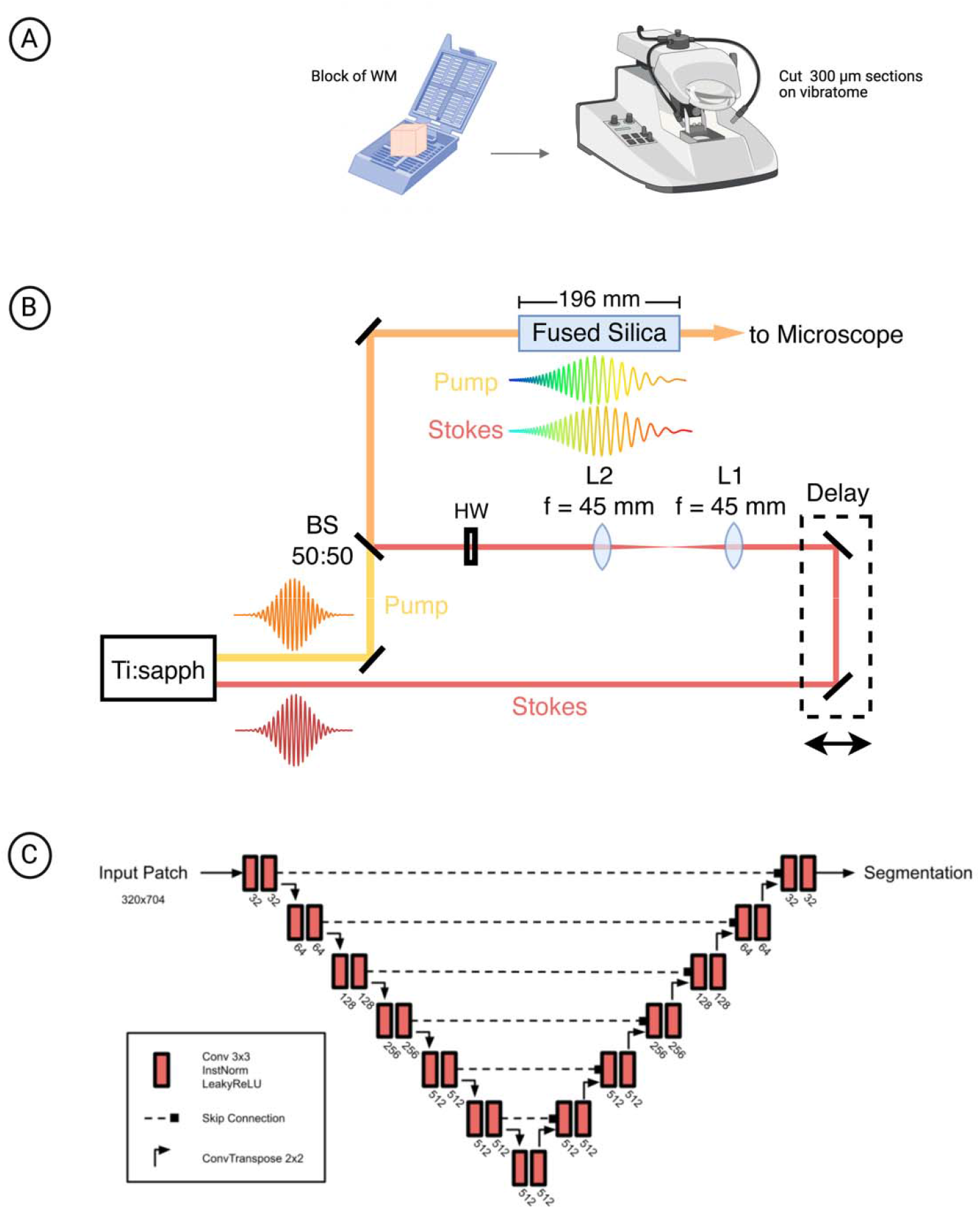
Summarized methods schematic. A) A block of white matter (approximately 1 cm^3^) is cut into 300 µm thick slices on a Leica Vibratome. B) sf-CARS acquisition setup. Short pulses of 100 fs from a Ti:sapphire (Ti:sapph) oscillator are outputted with a set polarization, frequency and power. Fixed-wavelength laser beam of 1045 nm (Stokes) is temporally delayed by a linear motion stage. It is then re-collimated through a 4f-system to match the collimation and size of the tunable (Pump) beam of 805 nm perfectly and then rotated with a half wave-plate (HW) to match the polarization of the pump. Both beams are split by a beamsplitter (BS) and subsequently synchronized and combined. They are then passed through a fused silica glass rod for a negative chirping effect or sliding of frequencies. Pulses are elongated, which improves the spectral resolution. The light is then sent to the microscope. C) Network architecture of the axon and myelin segmentation model. Each block contains convolution layers with 3×3 kernels, instance normalization and LeakyReLU. Skip connections merge the encoder outputs with the decoder inputs at each stage. The decoder applies 2D transposed convolutions to upsample the feature maps in between stages.

### sf-CARS microscopy

Sf-CARS imaging was performed using a Ti:Sapphire Insight X3 pulsed femtosecond (fs) laser system configured with two distinct output beams. The tunable laser was set to 805 nm with 100 fs pulses and ∼300 mW effective power at source output, while the fixed laser operated at 1045 nm with 100 fs pulses and ∼600 mW effective power at source ouput. Laser power settings were subsequently adjusted and optimized for each imaging session and sample type to achieve optimal contrast. Temporal chirping was achieved using a 19.6 cm fused silica rod with low dispersivity characteristics. Images were acquired through an Olympus 30X/0.8 NA water immersion objective with 2X additional optical zoom, utilizing a galvanometer and a polygonal mirror for scanning (Veilleux et al., 2008). The photomultiplier tube voltage was maintained at 750V, and each final image represented an average of 100 frames to enhance signal to noise ratio by reducing photon noise. Image acquisition (8 bits, maximum of 255) was performed over 500 × 1000 pixels at 30 frames per second. A tunable offset was used during the acquisition to eliminate low counts of photons and was adjusted for each image to optimize contrast and signal-to-noise ratio. Plus, both linear motion stages in the optical path (delay and at sample) were manually fine-tuned to achieve maximum resolution, and captured images underwent post-processing correction for enhanced contrast analysis. This sf-CARS acquisition setup is summarized in Figure 1B.

### AxonDeepSeg Deep Learning-based Morphometry Analysis

Deep learning-based axon segmentation typically comprises four sequential stages: data preparation, model training, result evaluation, and inference. During data preparation, microscopy images with corresponding axon and myelin annotations are normalized to consistent resolution parameters, partitioned into smaller image patches, and preprocessed using standardization and histogram equalization techniques. The model training phase utilizes manually segmented reference data to optimize network parameters. During inference, the trained model generates segmentation predictions on novel images. The model predictions are subsequently analyzed to derive morphometric measurements such as g-ratio, myelin thickness, axon diameter and eccentricity.

To quantify axon and myelin morphology within the UF, we use a deep learning-based segmentation and analysis pipeline integrated in AxonDeepSeg (Zaimi et al., 2018) itself based on our earlier work (Bégin et al., 2014). This approach facilitates automated, high-throughput analysis of the acquired CARS images. To optimize the quality of the semantic segmentation masks and enhance the robustness of the subsequent analyses, a custom model was trained specifically for this dataset, as detailed in the next section. Following segmentation of axonal and myelin structures, AxonDeepSeg automatically computed a comprehensive set of morphometric features for each identified axon. These features include axon diameter, myelin thickness, axon area and g-ratio. A *post hoc* screening procedure was implemented to filter out segmentation artifacts, such as axons with physiologically implausible g-ratios. This curated morphometric data was then utilized for subsequent statistical analyses and inter-group comparisons.

### Active Learning Segmentation Strategy

Axon and myelin segmentation was performed using a 2D U-Net architecture (“U-Net,” 2015) trained via an active learning strategy. This architecture is summarized in Figure 1C. This iterative approach enabled efficient model training with a progressively expanding dataset, concurrent with image acquisition, thereby maximizing segmentation accuracy while minimizing manual annotation effort. The model underwent eight iterative training cycles, each utilizing an expanded dataset. Initially, a model was trained using a small, manually annotated dataset comprising three 1000×500 images. For each subsequent iteration, the current model was applied to a new set of images from newly acquired subjects. Predictions selected for correction were chosen based on a qualitative assessment of segmentation quality, prioritizing those with the most apparent inaccuracies. Manual annotation was performed by four individuals with experience in neurohistological image interpretation. To ensure consistency, annotators were explicitly instructed to segment only those structures for which axonal identity could be definitively confirmed, excluding any ambiguous or uncertain cases from the training dataset.

These corrected segmentations were then incorporated into the training set, and the model was retrained. The number of annotators involved in manual correction increased to four as the active learning process progressed. Manual mask correction was performed in Gimp, Affinity Designer and Fiji. All 2D segmentation models were trained with the nnU-Net framework(Isensee et al., 2021). The final version of our dataset contained 124 images covering 6977 axons, and here we applied this final model to 6 “control” subjects. Our model can be used within the AxonDeepSeg software and is available for download at the following URL: https://github.com/axondeepseg/model-seg-human-brain-cars.

### Segmentation Post-Hoc Quality Control

Automated segmentation workflows can generate spurious artifacts that necessitate systematic filtering prior to morphometric analysis. To ensure data quality and remove false positive detections, a multi-criteria post-hoc screening procedure was implemented. Axons yielding physiologically implausible g-ratio values outside the range of 0 to 1 were excluded from analysis, as such values indicate erroneous segmentation of axonal or myelin boundaries. Additionally, axons with diameters below 0.335 μm were systematically removed from the dataset. Given the native imaging resolution of 0.166 μm per pixel, axons smaller than 0.335 μm correspond to structures with diameters less than 2 pixels, which are susceptible to inaccurate measurement due to partial volume effects and may represent either segmentation artifacts or genuine axonal structures below the reliable detection threshold of the imaging system. Additionally, axons exhibiting eccentricity values exceeding 0.9 were excluded from analysis, as the downstream quantification algorithm works under the assumption that axons have a circular shape, which may inadequately represent highly elliptical axonal profiles, potentially introducing systematic measurement errors. This quality control framework ensures that subsequent morphometric analyses are conducted exclusively on accurately segmented axonal structures with reliable geometric measurements.

Morphometric quantification of axon diameters was performed by calculating the diameter of an area-equivalent circle corresponding to each segmented axon. The g-ratio computation involves two diameter estimations: (1) inner diameter calculation utilizing solely pixels designated to the axon class, and (2) outer diameter calculation incorporating the aggregate of pixels assigned to both axon and myelin classes. This approach provides standardized measurements of myelination characteristics across the analyzed fiber population.

## Results

The subject information for the brain donors is shown in Table 1. After filtering for diameter greater than 0.335 and eccentricity less than 0.9, we were left with 2645 axons. An example segmentation is demonstrated in Figure 2. Axon diameters ranged from 0.37 µm to 6.38 µm, as demonstrated in the histogram in Figure 3A. The average axon diameter was 0.93 µm ± 0.54. We then stratified the axons by diameter bin, starting the cutoff at 0.335 µm to 0.35 µm, and then increasing by 0.25 µm increments until 1.75 µm. The final bin is 1.75 µm to the maximum diameter (1.75 µm +). The 0.5-0.75 µm bin contained the most axons, and the full binned distribution can be observed in Figure 3B.

**Subject information table.** Data are demonstrated as mean ± standard deviation.

|  |  |
| --- | --- |
| **N total** | 6 |
| **N male/female** | 3/3 |
| **Age** | 61.2 ± 17.8 |
| **pH** | 6.49 ± 0.34 |
| **PMI (h)** | 63.4 ± 20.2 |

**Figure 2.**
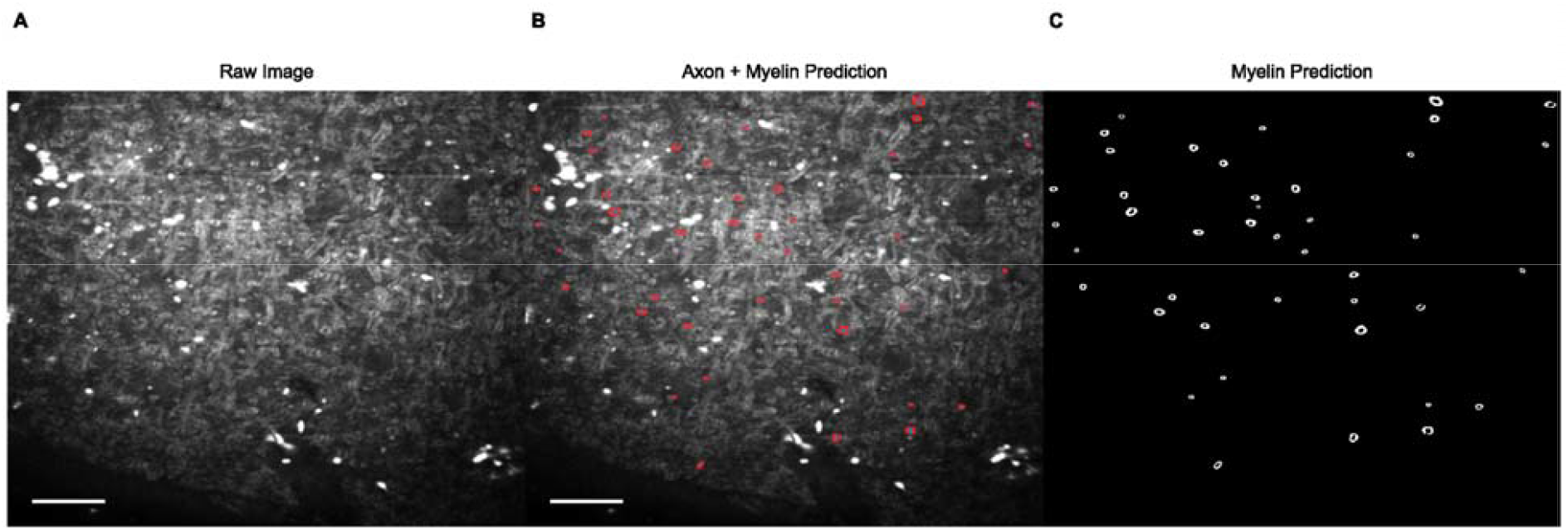
Example segmentation of axons and myelin from an sf-CARS image using custom trained AxonDeepSeg deep learning model. A) Raw, uncorrected sf-CARS image. Bright white spots indicate artifacts of concurrent 2-photon emission from tunable laser beam. B) Segmentation of myelin (red) and axon (blue) on top of raw sf-CARS image. C) Binary image of myelin segmentation. Scale bars = 20 µm

**Figure 3.**
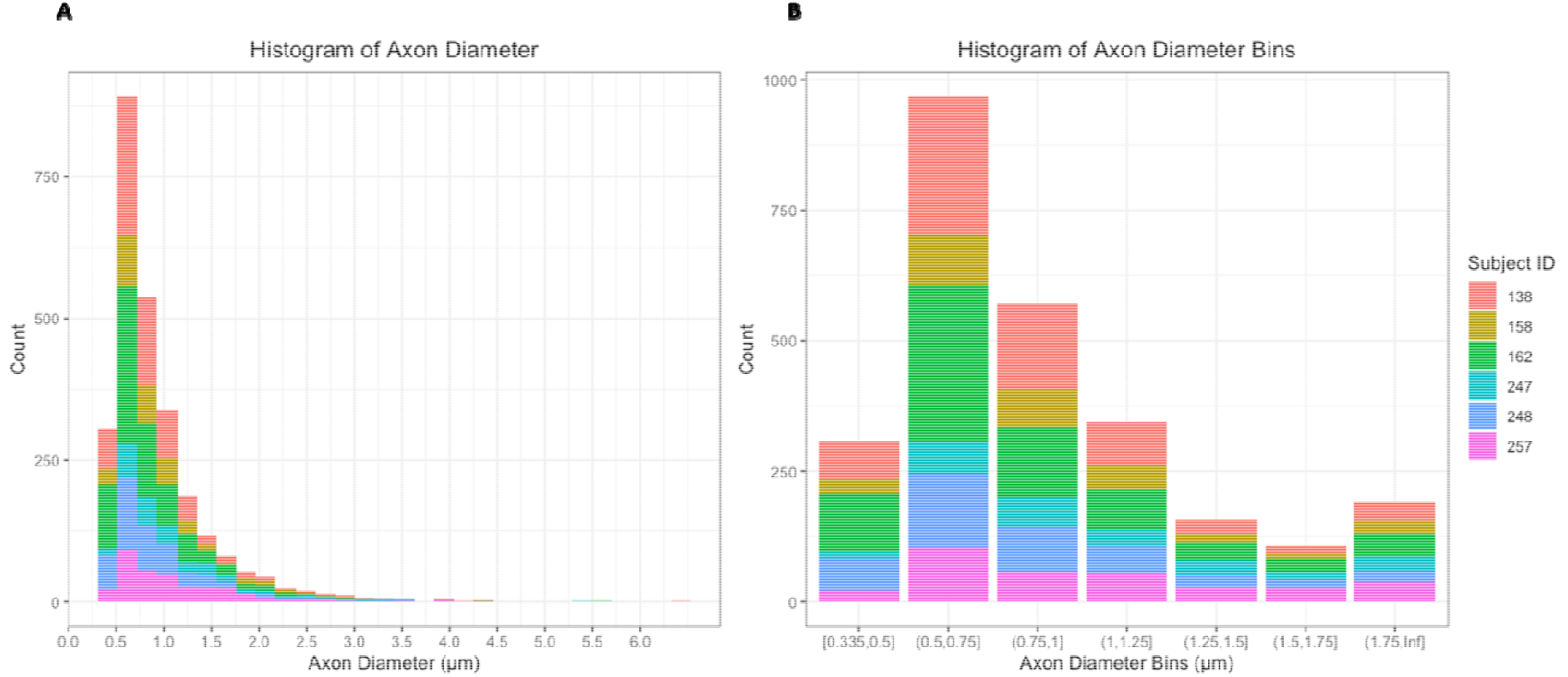
Axon diameter distributions. A) Histogram of axon diameter distribution, colored by individual subject ID. B) Histogram of binned axon diameters colored by individual subject ID.

Across all fibers, the mean myelin thickness = 0.48 µm ± 0.14, and the mean g-ratio = 0.47 µm ± 0.094. The mean myelinated fiber density is 844.36 axons/mm^2^. We cannot estimate the total axon density (which includes both myelinated and non-myelinated axons) because the sf-CARS is specifically tuned to map out lipid-rich structures like myelin. We validated that the morphometrics display the correct relationships with each other, summarized with a correlation plot (Figure 4A), with all correlations showing statistical significance (p < 0.001). All correlation coefficients were positive except for the correlation between myelin thickness and g-ratio, which are inversely correlated as g-ratio is measured as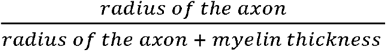. We observe that as expected, myelin thickness increases as a function of axon diameter (R^2^ = 0.38, p < 0.001) (Figure 4B). Similarly, as expected g-ratio increases as a function of axon diameter (R^2^ = 0.47, p < 0.001) (Figure 4C). The mean of each morphometric is summarized across axon diameter bins in Figure 5.

**Figure 4.**
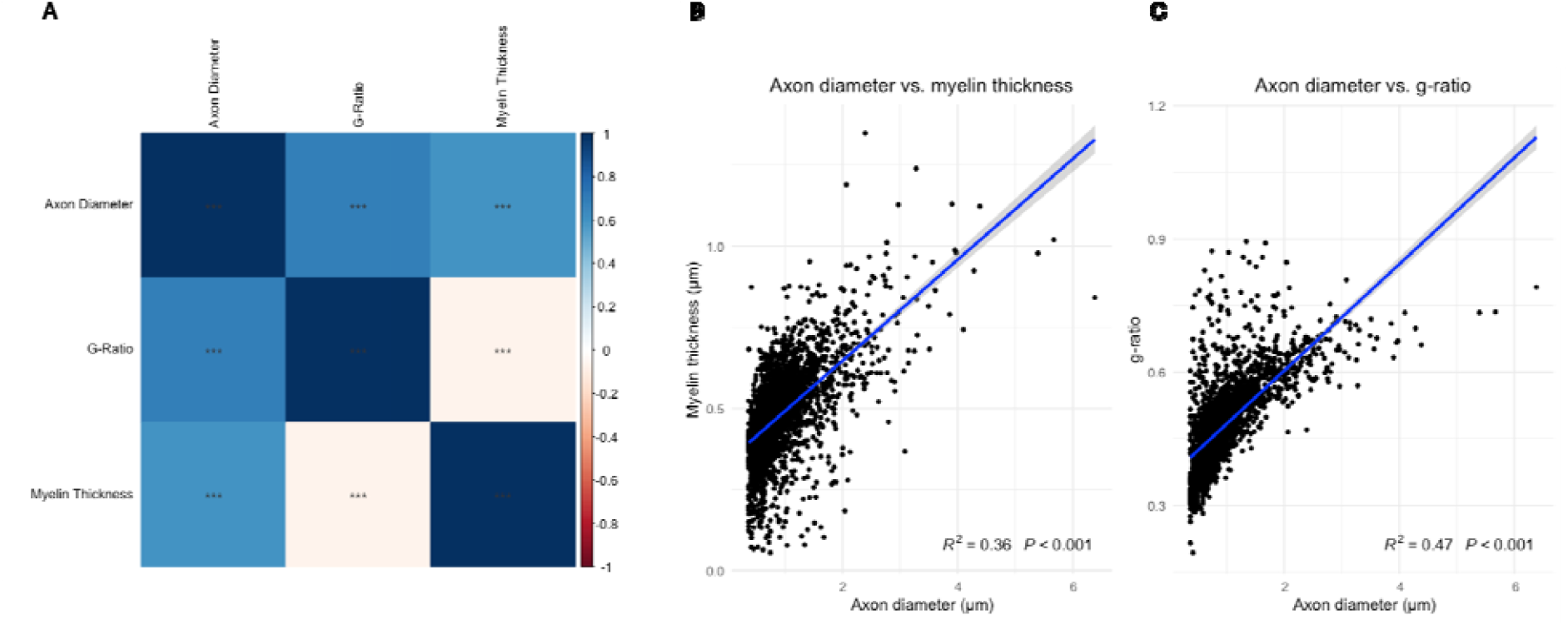
Morphometrics show the expected relationships. A) Correlation plot showing expected significant correlation across metrics. Color bar represents correlation coefficient with positive coefficients in blue and negative coefficients in red, and the significance stars represent p < 0.01. B) Scatterplot showing a directly proportional relationship between axon diameter and myelin thickness with trendline in blue (R^2^ = 0.38, p < 0.001) C) Scatterplot showing a directly proportional relationship between axon diameter and g-ratio with trendline in blue (R^2^ = 0.47, p < 0.001).

**Figure 5.**
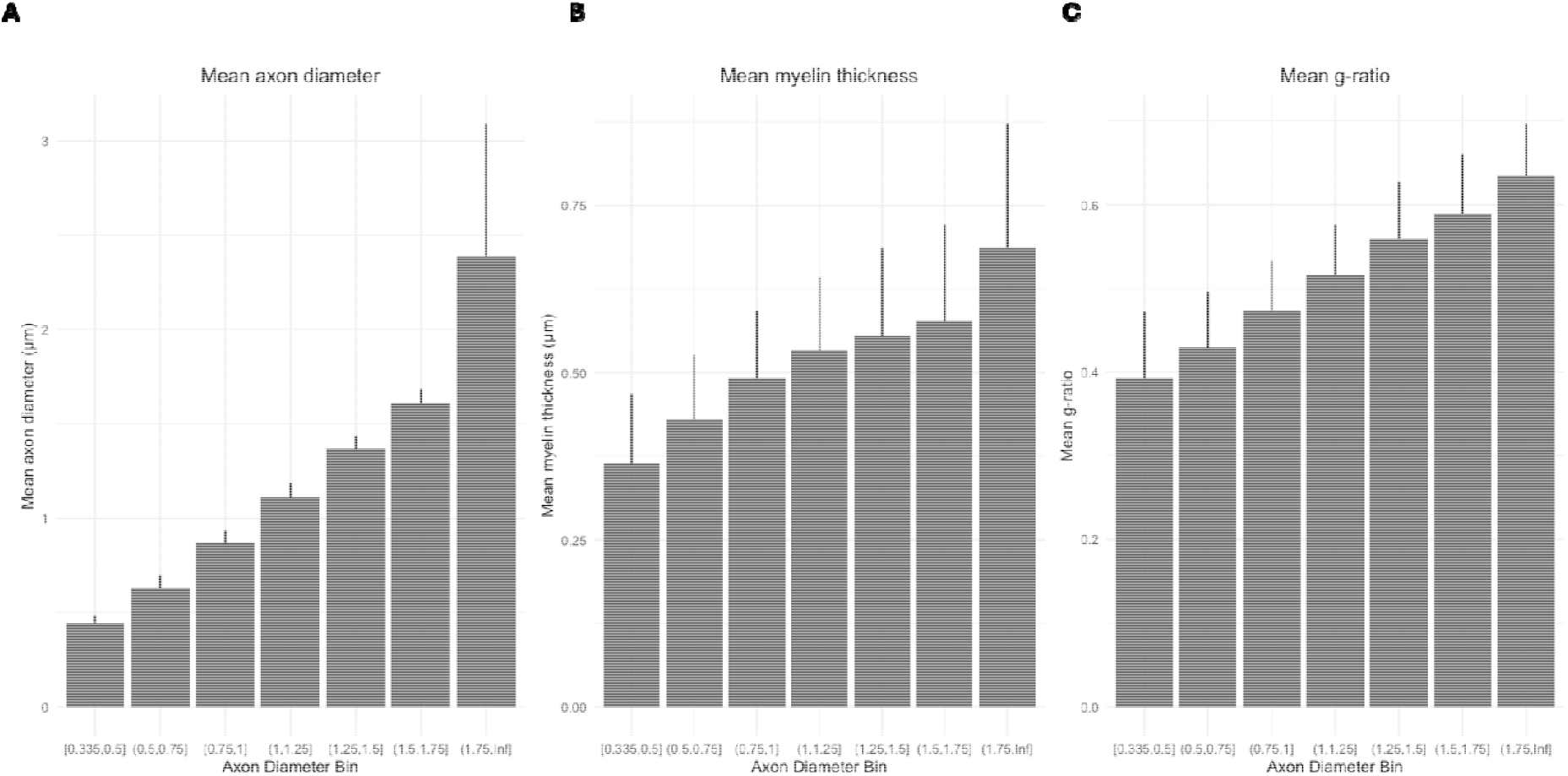
Bar plot demonstrating mean morphometrics across axon diameter bins. Data are shown as mean + standard deviation for A) axon diameter, B) myelin thickness, and C) g-ratio

To further validate the new pipeline, we elected to measure fibers of the ACC WM in a subject for which we also had UF fiber measurements (S247). We selected this region as it was used in Lutz et al., (2017), in which the previous picosecond-pulsed CARS system was employed. The ACC tissue was processed identically to the UF tissue. As such, we compared ACC fibers (n=206) and UF fibers (n=234) from the same subject, and observed that across all diameter bins, myelin was thicker, and g-ratio was smaller in the UF as compared to the ACC (Figure 6). The axon diameter and myelin thickness between regions appear similar across every diameter bin, except for those 1.75 µm+, in which the mean UF diameter is larger (Figure 6A).

**Figure 6.**
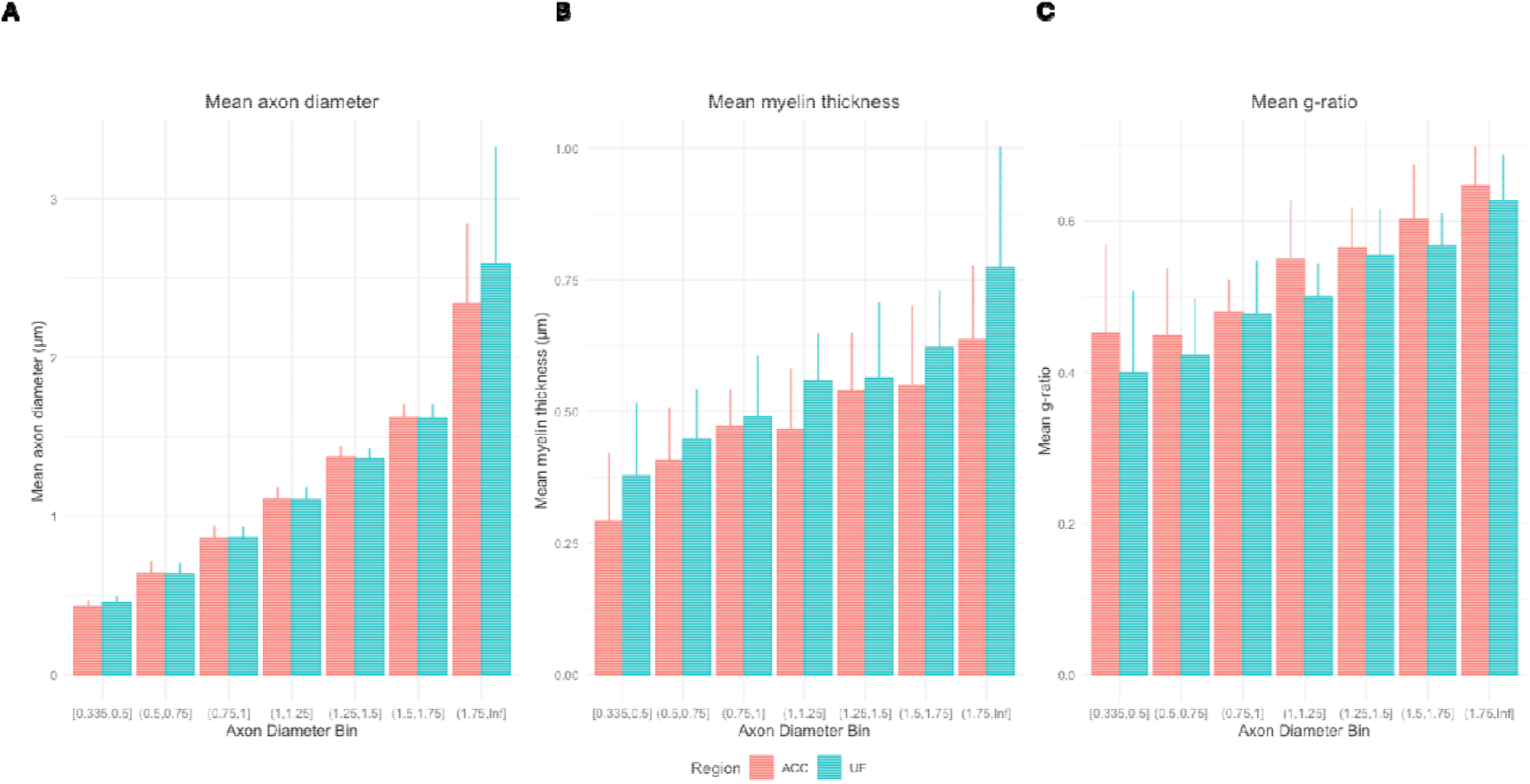
Bar plot demonstrating mean morphometrics for a single subject across regions stratified by axon diameter bins for A) axon diameter, B) myelin thickness, and C) g-ratio. Data for ACC is shown in pink and data for UF is shown in teal.

## Discussion

This article presents a full methodological pipeline for the automated ultrastructural analysis of human postmortem white matter. We validate the use of sf-CARS coupled with a custom trained AxonDeepSeg segmentation model for the quantification of the human postmortem UF. Furthermore, we provide the first ever characterization of UF ultrastructure, as well as a comparison between the ACC white matter and the UF. As expected, UF axons were larger and thicker than the ACC white matter axons, consistent with what is known about fiber bundles, which necessitate higher conduction velocities (Van Blooijs et al., 2023).

We also demonstrate that sf-CARS is feasible for use with human postmortem brain tissue. This approach, compared to the traditional picosecond-pulsed CARS systems used by Lutz et al. (2017), presents both significant advantages and inherent limitations. A primary strength lies in the utilization of more common and affordable Ti:sapphire laser technology, enhancing accessibility for research laboratories and allowing multimodal imaging in combination with 2-photon excitation fluorescence. The chirping mechanism substantially increases spectral resolution and enables sweeping of a broader Raman spectrum through larger chirped wavebands, facilitating more comprehensive and efficient information acquisition than previous systems (Pegoraro et al., 2009). This difference can be seen even with a low-dispersive rod, although the effect could be improved in further alterations. This approach also introduces the trade-off of dispersing laser power across the entire pulse spectrum, reducing effective power density at specific wavelengths. The current implementation remains suboptimal, as images still necessitate post-acquisition contrast correction, highlighting areas for technical refinement. While traditional picosecond systems may offer greater robustness (Evans et al., 2005), the sf-CARS method demonstrates superior adaptability and multiplicity of information capture. Future developments should focus on optimizing chirping efficiency through higher dispersity materials and exploring three-dimensional acquisition capabilities to enhance visualization of axonal structures and identify structural weaknesses at greater tissue depths. Additionally, this approach holds promise for monitoring myelin structural changes through differentiation and intensity analysis of the three distinct lipid peaks in Raman spectra, potentially providing novel insights into myelin pathology and neurodegeneration mechanisms.

Our data processing pipeline was better at detecting small diameter (<1 µm) axons compared to the tissue imaging and segmentation methods employed in Lutz et al. (2017). As such, it is key for the field to standardize the segmentation methods used, especially when making comparisons across different brain regions. The implementation of deep learning methodology enables unprecedented scalability in morphometric analysis, facilitating the quantification of substantially larger axonal populations than previously feasible with manual approaches (Oveisgharan et al., 2025; Tian et al., 2025). While automated segmentation inherently introduces potential sources of variability compared to expert manual annotation, our comprehensive post-hoc quality control protocol aims to mitigate these uncertainties, enhancing the reliability of extracted morphometric parameters across the analyzed dataset.

There are some noteworthy tissue-specific limitations to this study. The particular part of the UF used in this study (dissected at the temporal pole) represents the terminations of the fiber tract (Thiebaut De Schotten et al., 2012). In other words, the fibers tend to fan out to reach their targets, so the tract is therefore less densely compacted. As such, the fiber density is probably lower in this subsection compared to much of the UF. Since this analysis only considered the terminations of the UF temporal segment; these findings may not hold for the frontal segment or the insular segment (Ho et al., 2017). It would be beneficial for future studies to compare CARS ultrastructure at different segments of the UF to evaluate potential intra-tract variability in ultrastructure.

There are also important UF laterality differences reported (Olson et al., 2015; Von Der Heide et al., 2013) and, it is worth noting that we are only studying the left hemisphere in this study. For example, the left UF has been implicated in select supportive linguistic functions such as proper name retrieval (Papagno et al., 2011). On the other hand, psychopathy and acquired criminality studies show a consensus that the right UF shows lower fractional anisotropy values compared to controls (Kletenik et al., 2025). Other conditions report mixed findings with respect to laterality, such as those related to childhood abuse or socioemotional deprivation (Gur et al., 2019; Hanson et al., 2015). As such, it would be worthwhile to do a laterality comparison between the left and right UF, ideally at the temporal, insular, and frontal segments. Lastly, it is important to note that we cannot obtain all ultrastructural measures of interest with CARS, specifically pathology metrics that are only observable at very high resolution such as axonal swelling and myelin splitting.

In conclusion, we present and validate a full methodological pipeline for the specific detection, imaging, and automated segmentation of myelinated axons throughout postmortem brain sections to obtain quantitative ultrastructural metrics without requiring exogenous markers. We also provide the first ever descriptions of human UF ultrastructural metrics at the temporal segment. To ensure reproducibility, we supply detailed methodology on tissue fixation and slicing, sf-CARS imaging parameters, as well model training parameters and access to the trained deep learning model (https://github.com/axondeepseg/model-seg-human-brain-cars) which can be used within the existing AxonDeepSeg graphical user interface. We hope that researchers within the postmortem brain field will find this novel methodology both useful and easily reproducible. Future studies can use this pipeline to interrogate ultrastructural differences in neurological or psychiatric disorders in case-control studies.

## Acknowledgements

We would like to thank Sara-Maude Gagnon for her imaging assistance. Furthermore, we would like to thank the staff of the Douglas-Bell Canada Brain bank for the tissue dissections. Importantly, we would like to thank the brain donors and their next of kin who consented to donate the brains of their loved ones.

## Author contributions

KP: conceptualization, methodology, formal analysis, investigation, project administration, writing-original draft; JM: validation, investigation, data curation, writing - original draft; AC: conceptualization, methodology, software, validation, formal analysis, writing-original draft; VPN: conceptualization, methodology, validation, investigation, data curation, writing – review and editing; MM – methodology, software, validation;SJ: methodology; MAD: methodology, writing – review and editing; JCA: methodology, project administration, software, writing review and editing, supervision; DC: conceptualization, project administration, funding acquisition, writing review and editing, supervision; NM: conceptualization, project administration, funding acquisition, writing review and editing, supervision

